# An Electrophysiological Marker of Arousal Level in Humans

**DOI:** 10.1101/625210

**Authors:** Janna D. Lendner, Randolph F. Helfrich, Bryce A. Mander, Luis Romundstad, Jack J. Lin, Matthew P. Walker, Pal G. Larsson, Robert T. Knight

## Abstract

Deep non-rapid eye movement sleep (NREM) – also called slow wave sleep (SWS) – and general anesthesia are prominent states of reduced arousal linked to the occurrence of slow oscillations in the electroencephalogram (EEG). Rapid eye movement (REM) sleep, however, is also associated with a diminished arousal level, but is characterized by a desynchronized, ‘wake-like’ EEG. This observation challenges the notion of oscillations as the main physiological mediator of reduced arousal. Using intracranial and surface EEG recordings in four independent data sets, we establish the 1/f spectral slope as an electrophysiological marker that accurately delineates wakefulness from anesthesia, SWS and REM sleep. The spectral slope reflects the non-oscillatory, scale-free measure of neural activity and has been proposed to index the local balance between excitation and inhibition. Taken together, these findings reconcile the long-standing paradox of reduced arousal in both REM and NREM sleep and provide a common unifying physiological principle — a shift in local Excitation/ Inhibition balance — to explain states of reduced arousal such as sleep and anesthesia in humans.

**Significance Statement:** The clinical assessment of arousal levels in humans depends on subjective measures such as responsiveness to verbal commands. While non-rapid eye movement (NREM) sleep and general anesthesia share some electrophysiological markers, rapid eye movement sleep (REM) is characterized by a ‘wake-like’ electroencephalogram. Here, we demonstrate that non-oscillatory, scale-free electrical brain activity — recorded from both scalp electroencephalogram and intracranial recordings in humans — reliably tracks arousal levels during both NREM and REM sleep as well as under general anesthesia with propofol. Our findings suggest that non-oscillatory brain activity can be used effectively to monitor vigilance states.

## Introduction

Sleep and anesthesia both present with a behaviorally similar state of diminished arousal(1) and shared neurophysiologic features, namely increased low frequency power(2, 3) and a reduction in effective connectivity(4, 5). It has been argued that the reduced arousal in both states stems from a common neuronal mechanism. Current definitions of arousal vary and include e.g. autonomic, behavioral or mental arousal. An updated framework has been proposed recently(6). Here, we use the term arousal in its relation to vigilance states.

Most studies comparing sleep and anesthesia concentrated on slow-wave sleep and oscillatory dynamics such as slow waves (< 1.25 Hz)(1, 7, 8) as an increased activity in this frequency band has been associated with reduced arousal(1, 3). REM sleep is also associated with decreased arousal but is characterized by a desynchronized, active pattern in the electroencephalogram (EEG) similar to wakefulness(8). This paradox challenges the notion that changes in oscillatory activity such as slow waves are the exclusive determinant of reduced arousal.

Non-oscillatory, scale-free neural activity constitutes an important index of brain physiology and behavior(9–11). In the frequency domain, the scaling law between the power and the frequency of non-oscillatory brain activity can be estimated from the exponential decay of the power spectral density(9) and has previously been used to assess a variety of cognitive and EEG phenomena(12–18). A variety of terms have been used to describe this power-frequency relationship, such as power-law distribution, scale-free behavior, 1/f electrophysiological noise, fractal/spectral exponent(12, 17, 19) or fractal dynamics(9, 20–22). The exponent of the 1/f power-law distribution, also called spectral slope, differs between rest and task activity(9, 10) and changes with aging(21). Fractal dynamics and neural avalanches have also been observed in long-range temporal correlations of band-limited signals(23), however, it is likely that these two phenomena may reflect distinct entities with a different neurophysiological basis(9). Here, we focus on the fractal 1/f dynamics of the background activity.

Computational simulations indicate that the spectral slope provides a surrogate marker for the excitatory to inhibitory (E/I) balance with more negative slope values indexing enhanced inhibition(10, 20, 22) (Fig. S1), while others have observed the reversed pattern(11).

For this study, we followed the framework of Laureys et al. that defined consciousness on two axis – content (awareness) and level (arousal)(24). While the conscious content is low in NREM sleep and GABAergic anesthesia, it is high in wakefulness and dreaming states like REM. The arousal level, on the other hand, is low in all sleep states including REM. We hypothesized that states of reduced arousal are characterized by a shift of the E/I balance towards inhibition indexed by more negative slopes. To test this prediction, we analyzed four independent datasets: Electrophysiological recordings during sleep using either scalp EEG (Study 1, n = 20) or combined scalp and intracranial EEG (Study 2, n = 10; coverage see Fig. S2a) as well as under general anesthesia with propofol combined with scalp EEG (Study 3, n = 9) or intracranial EEG (Study 4, n = 12; subdural grid electrodes (electrocorticography; ECoG) and stereotactically placed depth electrodes (SEEG); coverage see Fig. S2b).

## Results

During a full night of sleep, the time-resolved spectral slope closely tracked the hypnogram (Fig. 1a). In the scalp EEG group (Study 1, n = 20; a baseline rest recording was available in n = 14), we observed a decrease from values of −1.87 ± 0.18 (mean ± SEM) during quiescent rest to −3.46 ± 0.16 in NREM (N3) and −4.73 ± 0.23 in REM sleep (Fig. 1b). These differences were significant across all scalp EEG channels (repeated-measures ANOVA: p < 0.0001, F_1.94, 25.17_ = 56.05, d_Rest-Sleep_ = 3.07). Furthermore, N2 sleep exhibited an average slope of −3.67 ± 0.10 that was also significantly below rest (n = 14; p_Rest-N2_ < 0.0001; t_13_ = 7.97; d_Rest-N2_ = 3.31; Fig. S3a). Post-hoc t-tests (uncorrected) revealed a significant difference between rest and N3 (p_Rest-N3_ < 0.0001, t_13_ = 5.69, d_Rest-N3_ = 2.49), between rest and REM (p_Rest-REM_ < 0.0001, t_13_ = 11.67, d_Rest-REM_ = 3.71) and between N3 and REM sleep (p_N3-REM_ = 0.0007, t_13_ = 4.44, d_N3-REM_ = 1.70).

**Fig. 1:**
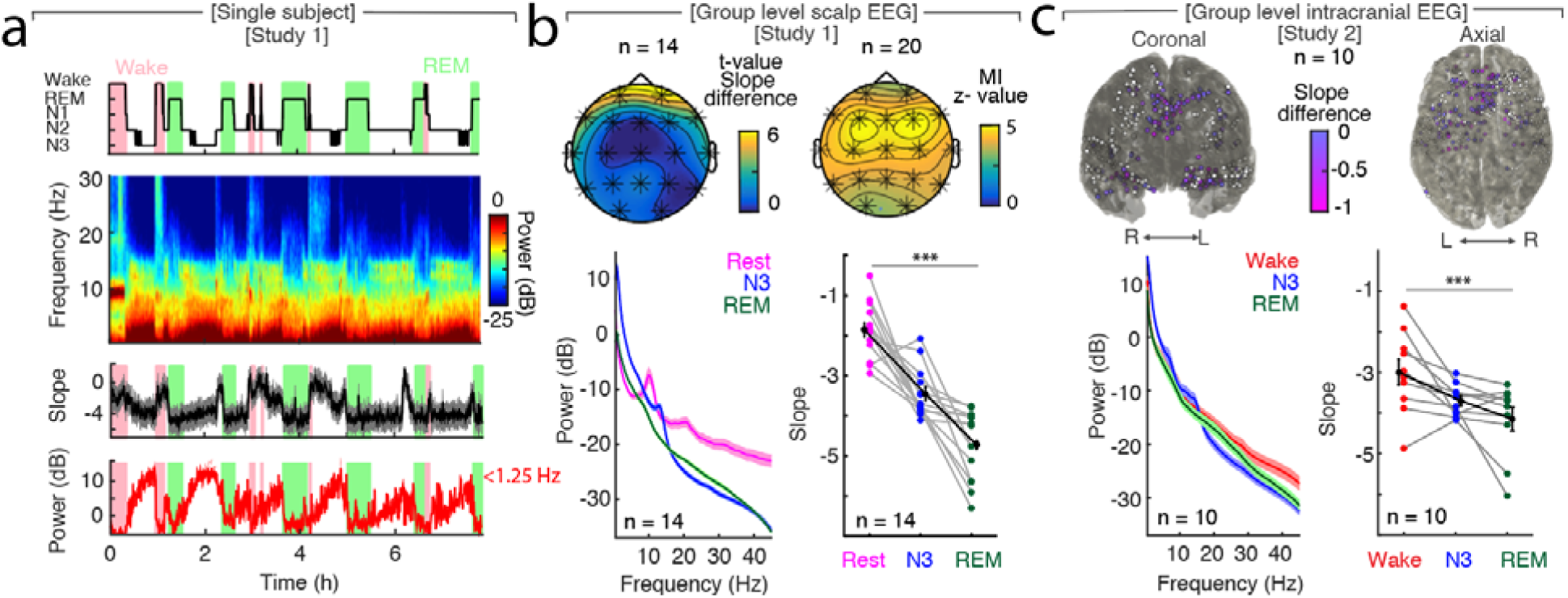
The spectral slopes tracks changes of arousal level in sleep. **a**, Time-resolved average of three frontal EEG channels (F3, Fz, F4) during a night of sleep. Upper panel: Expert-scored hypnogram (black), wake (pink), REM (light green). Upper middle: Time-frequency decomposition. Lower middle: Spectral slope (black; mean ± SEM). Lower panel: Low-frequency (<1.25 Hz) power (red; mean ± SEM). **b**, Sleep in scalp EEG. Upper panel: Left: Cluster permutation test of slope difference between sleep and rest (n = 14). * p < 0.05. Right: Mutual Information between the time-resolved slope and hypnogram (n = 20). Cluster permutation test against surrogate distribution created by random block swapping: * p < 0.05. Lower panel: Left - Power spectra (n = 14; mean ± SEM); Right – Spectral slope (n = 14). Rest (magenta), NREM stage 3 (blue), REM sleep (green) and grand average (black; mean ± SEM). Repeated measures ANOVA: *** p < 0.001. **c**, Sleep in intracranial study (n = 10). Upper panel: Left – coronal, right – axial view of intracranial channels that followed (magenta) or did not follow (white) the EEG pattern of a lower slope during sleep (REM/NREM 3). Lower panel: Left – Power spectra (mean ± SEM); Right – Spectral slope of simultaneous EEG recordings (Fz, Cz, C3, C4, Oz). Wakefulness (red), NREM stage 3 (N3; blue), REM sleep (green) and grand average (black; mean ± SEM). Repeated measures ANOVA: *** p = 0.001.

If all the available wake periods before, during and after the sleep recordings were utilized for slope analysis (n = 20), it resulted in a higher variability across subjects during wakefulness (Fig. S3b), which can be explained by the fact that the subjects were already or still drowsy and data during state transitions was included. However, the overall pattern was remarkably similar (Fig. S3c). As this approach increased our available data, we used all wake trials (referred to as wake) for subsequent analysis.

To assess where on the scalp the slope tracks arousal states best, we calculated the Mutual Information (MI) between the time-resolved spectral slope and the hypnogram in all 20 subjects. We observed a significant positive cluster across all sensors, which peaked over frontal electrodes F3, Fz and F4 (Fig. 1b).

Cranial muscle activity has similar frequency characteristics in the 30-70 Hz range and might confound spectral slope estimates. Therefore, we controlled for any impact of muscle activity by repeating the analysis after local referencing (Laplacian, p_Spearman_ < 0.001; p_MI_ < 0.0001) and additionally utilized partial correlations that considered the slope of the electromyography (EMG) as a confounding variable (p_Spearman_ < 0.001). All control analyses confirmed that the observed effect was not confounded by muscle activity (Fig. S4).

During REM sleep, power in the slow wave range (SO power; <1.25 Hz) was comparable to wakefulness corroborating the observation of a ‘wake-like’ EEG pattern in REM and the paucity of slow oscillations (p = 0.423, t_18_= −0.82, d = −0.25; Fig. 1a). The spectral slope, however, was significantly different between these states. To further quantify this effect, we trained a classifier (linear discriminant analysis; LDA) to discriminate between REM sleep and wakefulness using either the spectral slope or SO power (n = 18). The classifier performance was significantly better for the spectral slope compared to SO power when differentiating between REM and waking (78.75 ± 2.98 % (mean ± SEM) vs. 60.03 ± 3.72 %; p = 0.0023, t_17_ = 3.58, d_Slope-SO power_= 1.21, chance level: 50 %). When differentiating between N3 sleep and wakefulness, both spectral slope and SO power had a classifier performance that was significantly above the 50 % chance level (for slope p < 0.001 vs. for SO power p < 0.001) and comparable to each other (73.05 ± 2.97 % for spectral slope vs. 82.09 ± 2.13 % for SO power, p = 0.0423, t_17_ = −2.19, d_Slope-SO power_ = −0.83). Likewise, when all three states were classified simultaneously, both SO power and the spectral slope performed well above chance (chance = 33%; SO: 64.94 ± 2.04%, mean ± SEM; t_17_ = 15.04, p < 0.001, d = 5.01; slope: 58.09 ± 2.35%; t_17_ = 10.55, p < 0.001, d = 3.52) and did not differ in the overall performance (t_17_ = −1.80, p = 0.0899, d = −0.63). This is due to the fact that SO power is advantageous to classify N3 sleep, while the slope is superior to detect REM sleep. Notably, significant classification is also possible when the spectral slope is estimated at lower frequencies (e.g. 1-20 Hz; 84.19% ± 2.46, paired t-test vs. chance (33%): p < 0.001, t_17_ = 20.64, d = 6.88). This effect is partly driven by an increase in low frequency power needed to correctly classify N3, and is equivalent to using SO power, but the 1-20 Hz ranges does not track wakefulness and REM, thus, reducing mutual information with the hypnogram (see also Fig. S7).

These results reveal that the spectral slope is a more powerful predictor of REM sleep than SO power and also reliably discriminates deep N3 sleep from wakefulness. Furthermore, classification based on the spectral slope provides comparable accuracy levels in discriminating REM from wakefulness as trained personnel, given that the inter-rater reliability between sleep scoring experts is typically about 80%(25). Finally, the discrimination between REM and waking using the spectral slope does not require simultaneous electrooculography (EOG) or EMG recordings but can be detected solely from the electrophysiological brain state.

In the intracranial recording group (Study 2, n = 10), the simultaneous EEG recordings (Fz, Cz, C3, C4, Oz) again displayed a more negative spectral slope for reduced arousal levels: From −2.99 ± 0.32 (mean ± SEM) in wakefulness the slope decreased to −3.69 ± 0.12 in NREM (N3) to −4.15 ± 0.29 in REM sleep (Fig. 1c). Again, these three states were significantly different in a repeated-measures ANOVA (p = 0.001; F_1.97, 17.74_ = 10.79, d_Wake-Sleep_ = 1.12). Post-hoc t-tests (uncorrected) showed a significant difference between wakefulness and REM (p < 0.001; t_9_ = 4.78; d = 1.19) and wakefulness and N3 (p = 0.026; t_9_ = 2.66; d = 0.97) but not between N3 and REM (p = 0.098; t_9_ = 1.84; d = 0.64).

The intracranial SEEG contacts that mirrored the observed scalp EEG pattern (more negative spectral slope in N3 and REM; 155 of 352 SEEG (44.03 %; significantly above chance; *X*^2^ = 8.20, p = 0.0042; chi-squared test); Fig. 1c) exhibited a clear anatomical distribution centered in the medial prefrontal cortex and medial temporal lobe structures (for Wake - N3 and Wake - REM see Fig. S5a, b; grid electrodes see Fig. S6a, b), hence, converging on the very same brain regions known to be the most relevant for sleep-dependent memory consolidation(26–29). Note that we did not specifically target any brain regions and in contrast to previous studies using grid electrodes(9, 22), the majority of our probes were stereo-tactically placed depth electrodes. Given the spatial heterogeneity of intracranial responses(30), the convergence on medial PFC nicely resembles the observed scalp pattern as observed at the overlying scalp EEG electrode Fz. The distinct intracranial spatial pattern combined with the bipolar referencing scheme again confirms that the results are not confounded by muscle activity.

To verify the chosen fit parameters, we reanalyzed the correlation and MI analysis between hypnogram and time-resolved slope as a function of different center frequencies and window lengths (Fig. S7). In addition, we explored a wide-range of fit parameters after discounting the oscillatory components from the PSD by means of irregular resampling (IRASA(26, 31)). All control analyses corroborated our findings and indicate that the spectral slope in the range from 30 - 45 Hz reliably tracks arousal levels and behavioral state transitions (Fig. S7). Given the relationship between spectral edge frequency and median frequency(32), we assessed the relationship between SO power and the PSD slope. We observed that the SO power explains 7.9 ± 0.01% (mean ± SEM) of the variance in the slope, however, a partial correlation with SO power as a confound does not change the correlation between slope and hypnogram (Fig. S7).

On the scalp level, the trough of a slow wave is associated with cortical ‘down-state’, while the peak reflects an ‘up-state’(33, 34). The spectral slope was able to reflect these rapid changes during sleep with a more negative 1/f slope observed at troughs compared to peaks (Fig. S8). This effect was most pronounced over frontal channels (cluster-based permutation test: p = 0.005, d_Trough-Peak_ = −0.65).

Slow waves are detected in slow-wave sleep but are also observed during REM sleep(35) as well as wakefulness(36); albeit less prevalently. We detected a significantly higher number of slow waves during N3 sleep (SO_N3_ = 28.79 ± 0.79 per minute; mean ± SEM) compared to REM sleep (SO_REM_ = 2.16 ± 0.89 per minute; SO_N3-REM_: p < 0.0001, t_19_ = 22.64, d = 7.05) and wakefulness (SO_Wake_ = 5.05 ± 0.51 per minute; SO_N3-Wake_: p < 0.0001, t_19_ = 25.32, d = 6.92; Extended Data Fig. 9c). Interestingly, the averaged slope at the through of the slow waves was significantly different between arousal states: −3.40 ± 0.09 in slow-wave, −4.00 ± 0.18 in REM sleep and −2.26 ± 0.12 in wakefulness (mean ± SEM) mirroring our observation of the overall slope differences (Fig. S9c; uncorrected for multiple testing: Wake-N3: p < 0.0001, t_18_ = 7.07, d = 2.38; Wake-REM: p < 0.0001, t_18_ = 9.67, d = 2.55, N3-REM: p = 0.01, t_19_ = 2.73, d = 0.91). Therefore, the spectral slope is able to discern arousal even during slow wave events.

To test if the state-dependent modulation of the spectral slope was sleep-specific or generalized to other forms of decreased arousal, we analyzed two datasets obtained during general anesthesia with propofol. Under propofol anesthesia the time-resolved spectral slope again closely tracked changes in arousal level (Fig. 2a). In both scalp and intracranial EEG, we observed a more negative spectral slope under anesthesia compared to wakefulness (Fig. 2b, c): In the scalp EEG group (Study 3, n = 9), we found a decrease from −1.81 ± 0.29 (mean ± SEM) during wakefulness to −3.10 ± 0.19 under anesthesia. This difference was significant (paired t-test: p < 0.0001, t_8_ = 7.73, d_Wake-Anesthesia_ = 1.71) and in a cluster-based permutation test, the effect formed one single cluster spanning all 25 electrodes (p < 0.001).

**Fig. 2:**
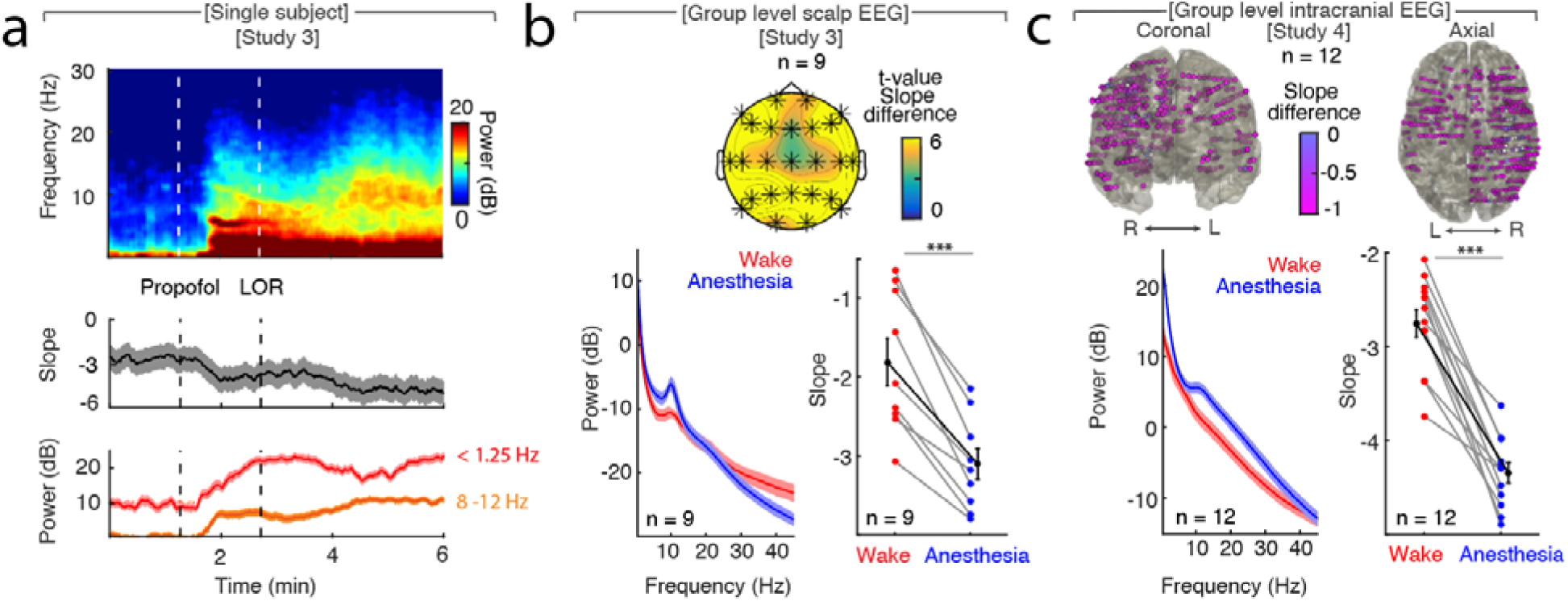
The spectral slope tracks changes in arousal level under general anesthesia with propofol. **a,** Time-resolved average of 35 intracranial frontal channels during anesthesia. Upper panel: Time-frequency decomposition. Dotted white lines: Induction with propofol, loss of responsiveness (LOR). Middle: Spectral slope (black; mean ± SEM). Lower panel: Low frequency (<1.25 Hz; red) and alpha (8 – 12 Hz; orange) power (mean ± SEM). **b**, Anesthesia in scalp EEG (n = 9). Upper panel: Spatial extent of spectral slope difference. Cluster permutation test: * p < 0.05. Lower panel: Left - Power spectra (mean ± SEM); Right – Spectral slope. Wakefulness (red), anesthesia (blue) and grand average (black; all mean ± SEM). Paired t-test *** p < 0.001. **c**, Anesthesia in intracranial recordings (n = 12). Upper panel: Left – coronal, right – axial view of slope difference. Lower panel: Left – Power spectra; Right – Spectral slope. Wakefulness (red), anesthesia (blue) and grand average (black; mean ± SEM). Paired t-test: *** p < 0.001.

In the intracranial recordings (Study 4, n = 12), we observed a spectral slope of −2.75 ± 0.15 during wakefulness and −4.34 ± 0.11 under anesthesia. Again, this difference was significant (paired t-test: p < 0.0001, t_11_ = 9.93, d_Wake-Anesthesia_ = 3.57) and could be detected in the majority of electrodes (470 of 485 SEEG (96.9 %); Fig. 2c). Patients who were implanted with surface grid in addition to depth electrodes (n = 4) showed the same pattern: The spectral slope decreased from wakefulness to anesthesia in the majority of the recording sites (129 of 147 ECoG (87.75 %); Fig. S6c).

These findings demonstrate that the spectral slope reliably differentiates between wakefulness and general anesthesia in humans(22). Future studies will be needed to determine the reliability of this marker on larger cohorts to establish clinical usability. In both scalp and intracranial recordings, we observed a brain-wide decrease in the spectral slope, supporting the notion that propofol anesthesia induces a global brain-wide state of increased inhibition(8).

## Discussion

Collectively, the results from these four studies provide five main advances. First, the spectral slope tracks changes in arousal levels in both sleep and anesthesia with high temporal precision from sub-second epochs to full night recordings. Note that the slope differences between wakefulness and states of reduced arousal show a similar pattern on the scalp level (Fig. 1b, 2b).

According to the framework proposed by Laureys et al., consciousness can be assessed on two axis – the content (e.g. awareness) and the level (e.g. arousal)(24), however, an updated framework has recently been proposed(6). Our definition of arousal is similar to what has been described as vigilance. However, our neurophysiological investigations did not set out to test one specific framework, but we do interpret our findings in light of previously published definitions. Hence, we assume that our marker does not track conscious thoughts, content or awareness, but indexes a vigilance state. While the arousal level is reduced in all three states, conscious content is thought to fluctuate during sleep, mostly in the form of dreams during REM(37). Thus, measures such as the Perturbational Complexity Index(38) that might track the level of consciousness are decreased in slow-wave sleep and GABAergic anesthesia but are maintained to a certain degree during REM sleep and ketamine anesthesia, both states associated with vivid dreams(37–39). These measures are unable – unlike the spectral slope – to reliably differentiate arousal levels, e.g. wakefulness and REM. Previous studies in rodents identified markers of reduced arousal in sleep and under general anesthesia, namely fronto-parietal theta and high-gamma connectivity(39, 40). In several control analyses we found that the spectral slope was superior to fronto-parietal theta connectivity in tracking sleep stage dependent dynamics (p < 0.0001, t_19_ = 7.01; d = 2.22) and in reliably differentiating REM and slow wave sleep (Fig. S10). Our dataset did not have a sufficient number of electrodes in the parietal lobe to extend the analysis to the high-gamma band since this is an infrequent site for epilepsy.

Second, the spectral slope provides a mechanistic explanation – a shift of the E/I balance towards inhibition – for the reduced arousal level in both slow-wave and REM sleep. The estimation of local E/I balance has been limited to invasive single cell recordings with a classification of neuron subtypes into excitatory and inhibitory cells(22). Recent computational simulations, however, demonstrated that local E/I balance can be inferred from changes of the spectral slope: An increase in inhibition results in a decrease of slope(11, 22). Our results of a decreased slope in slow-wave and REM sleep as well as under general anesthesia may be explained by an increase in inhibition. This interpretation is supported by results of single cell studies in animals that reported a reduction of multiunit or pyramidal cell activity during not only in slow-wave but also in REM sleep(41–44). Interestingly, REM exhibited a significantly lower slope than slow-wave sleep (Fig. S11). This result is in line with previous studies reporting a lower neuronal firing rate for REM sleep compared to slow-wave sleep(41, 43, 44) that was associated with an increase in inhibitory activity(41, 43). While these lines of research converge on the notion that the spectral slope tracks the E/I balance of the underlying population, it might also reflect changes in firing rate or synchronization. A testable hypothesis that arises from our observations is that cell-type specific causal manipulations by optogenetics (e.g. pyramidal and SOM interneurons) should bias the spectral slope in opposite directions.

Previous studies utilized a variety of different fit parameters and it is currently unclear what the ‘best’ range for slope fitting is(12–18). It had been suggested that fits to different frequencies might index different properties of the underlying population activity(9, 14, 16, 17). Our results that demonstrate that the range from 30-45 Hz best correlates (and exhibits significant mutual information) with the hypnogram, which is in line with recent modeling work indicating a similar range(22). Future studies involving single neuron recordings will be needed to unravel the precise relationship between population firing statistics and band-limited changes in the PSD slope. We believe that in particular comparative studies involving rodents(22, 45), primates(22) and humans combined with modeling work has the potential to integrate divergent findings into a coherent framework and to determine the neurophysiologic basis of the spectral slope. It will be of substantial interest to assess whether neurophysiological mechanisms are preserved across species, which greatly vary in anatomy, in particular in the prefrontal cortex(46, 47).

Third, the rapid changes in spectral slope observed over the course of a slow wave are in accordance with the notion that these oscillations orchestrate cortical activity during sleep by interleaving periods of neural silence with enhanced neural activity(41). This suggests that E/I balance and arousal level during slow wave sleep are not constant but wax and wane on a short time scale – whereas they seem to be more constant during REM sleep(41). This finding is in line with the active, maximal inhibition during REM sleep observed in single cell recordings of animal cortices(41, 43) and could explain why epileptic seizures during the night occur predominantly in NREM and rarely during REM sleep(48).

Fourth, our observations support the premise that anesthesia is a brain-wide state(8), whereas sleep exhibits network-specific activity patterns (e.g. between the PFC and the hippocampus)(49). This is especially relevant considering the theories of active memory processing in sleep(50, 51).

Fifth, the spectral slope can be reliably estimated from scalp EEG recordings, providing a potential tool that can be incorporated into intraoperative neuromonitoring, automatic sleep stage classification algorithms and tracking other states of reduced arousal such as epileptic seizures, coma and the vegetative or minimally conscious state.

## Data Availability

Data generated and/or analyzed in the current study is available from the corresponding author upon reasonable request.

## Code availability

Custom code used for analyzing the datasets of the current study is available from the corresponding author upon reasonable request.

## Acknowledgement

This work was supported by Grant LE 3863/2-1 of the German Research Foundation (Deutsche Forschungsgemeinschaft (J.D.L.), a National Institute of Neurological Disorders and Stroke Grant R37NS21135 (R.T.K., J.D.L.), the Alexander von Humboldt Foundation (Feodor Lynen Program; R.F.H.), an intramural fellowship from the Dept. of Psychology, University of Oslo (R.F.H.), R01AG03116408 (M.P.W.), RF1AG05401901 (M.P.W.), RF1AG05410601 (M.P.W.) and F32-AG039170 (B.A.M.), all from the National Institute of Health. We thank Jie Zheng, Julia Kam and the EEG technicians at UC Irvine Medical Center for their assistance and all the patients for their participation.

## Author Contribution

Conceptualization, J.D.L, R.F.H., M.P.W. and R.T.K.; Methodology, J.D.L. and R.F.H.; Software, J.D.L. and R.F.H.; Validation, J.D.L. and R.F.H.; Formal Analysis, J.D.L. and R.F.H.; Investigation, J.J.L., B.A.M., L.R. and P.G.L.; Resources, P.G.L, J.J.L., M.P.W. and R.T.K.; Data Curation, J.D.L, R.F.H., J.J.L., P.G.L., B.A.M., M.P.W. and R.T.K.; Writing – Original Draft, J.D.L.; Writing – Review & Editing, J.D.L., R.F.H., P.G.L, B.A.M., M.P.W. and R.T.K.; Visualization, J.D.L. and R.F.H.; Supervision, R.T.K.; Project Administration, P.G.L., J.J.L., M.P.W. and R.T.K.; Funding Acquisition, R.T.K.

## Materials and Methods

### Participants

We collected four independent datasets for this study to assess the neurophysiological basis of states of reduced arousal, namely sleep and general anesthesia. We recorded either non-invasive scalp electroencephalography (EEG) or intracranial EEG (electrocorticography; ECoG) using surface grid and strip electrodes and stereotactically placed depth electrodes (SEEG; for coverage see Fig. S2).

### Sleep

#### Study 1 - Sleep scalp EEG

Study 1 was conducted at the University of California at Berkeley. All participants were informed and provided written consent in accordance with the local ethics committee (Berkeley Committee for Protection of Human Subjects Protocol Number 2010-01-595). We analyzed recordings from 20 young healthy participants (20.4 ± 2.0 years, mean ± SD; 12 female). Polysomnography was recorded during over an 8-hour period as well as during 5 min quiescent rest with eyes closed before and after sleep. Data was recorded on a Grass Technologies Comet XL system (Astro-Med, Inc., West Warwick, RI) with a 19-channel EEG using the standard 10-20 setup as well as three electromyography (EMG) and four electro-oculography (EOG) electrodes at the outer canthi. The EEG was referenced to the bilateral linked mastoids and digitized at 400 Hz (0.1 to 100 Hz)(26, 27, 52, 53). Sleep staging was carried out by trained personnel and according to the newest guidelines(54).

#### Study 2 - Sleep intracranial EEG

Study 2 was conducted at the University of California at Irvine, Medical Center. Ten epilepsy patients (6 female) undergoing invasive pre-surgical localization of their seizure focus were included in this study. All patients provided informed consent according to the local ethics committees of the University of California at Berkeley and at Irvine (University of California at Berkeley Committee for the Protection of Human Subjects Protocol Number 2010-01-520; University of California at Irvine Institutional Review Board Protocol Number 2014-1522, UCB relies on UCI Reliance Number 1817) and gave their written consent before data collection. They were between 22 and 55 years old (33.1 ± 11.5 years; mean ± SD). Electrode placement was solely dictated by clinical criteria (Ad-Tech, SEEG: 5 mm inter-electrode spacing; Integra, Grids: 1 cm, 5 or 4 mm spacing). Data was recorded with a Nihon Kohden recording system (256 channel amplifier, model JE120A), analogue-filtered above 0.01 Hz and digitally sampled at 5 kHz. To facilitate gold-standard sleep staging, simultaneous EOG, electrocardiography (ECG) from 5 leads and EEG was recorded by exemplary electrodes of the 10 - 20 setup depending on the localization of the intracranial electrodes but mostly consisting of Fz, Cz, C3, C4 and Oz. A surrogate EMG signal was derived from the ECG and EEG by high-pass filtering above 40 Hz. Sleep staging was carried out by trained personnel.

### Anesthesia

The EEG and intracranial anesthesia studies were conducted at the University Hospital of Oslo. All participants or their parents provided informed written consent according to the local ethics committee guidelines (Regional Committees for Medical and Health Research Ethics in Oslo case number 2012/2015 and extension 2012/2015-8) and the Declaration of Helsinki.

#### Study 3 - Anesthesia scalp EEG

Ten patients (2 female) undergoing anterior cervical discectomy and fusion participated in Study 3 and received a total intravenous anesthesia with remifentanil and propofol. They had an American Society of Anesthesia status of I - III, were between 46 and 64 years old (53.3 ± 5.7 years; mean ± SD) and otherwise healthy. Data was recorded from the induction of anesthesia to the recovery from 25 channel EEG according to the 10 - 20 layout (EEG Amplifier, Pleasanton, California, USA) with an additional row of electrodes (F9, F10, T9, T10, P9, P10) at a digitization rate of 512 Hz, or in the case of one patient at 256 Hz. The electrode for referencing was placed at CP1. Three patients were not recorded for the planned entire time span – one recording was only started after induction, while two were stopped before recovery(55).

#### Study 4 - Anesthesia intracranial EEG

A total of 12 patients (3 female) with intractable epilepsy participated in Study 4. They were between 8 and 52 years old (26.6 ± 13.2 years; mean ± SD). Data was collected during the explantation of the intracranial electrodes from induction of anesthesia up to the point of their removal. All patients received total intravenous anesthesia with propofol and remifentanil at the University Hospital of Oslo. All patients were placed back on their usual antiepileptic medication before the procedure. Data was recorded on a Natus NicoletOne system with a 128-channel capacity and a digitization rate of 1024 Hz for up to 64 or 512 Hz for up to 128 channels.

### Anesthetic management

All patients received a premedication with 3.75 to 7.5 mg midazolam (Dormicum®, Basel, Switzerland); the anesthesia scalp EEG group (Study 1) received additional 1 g oral paracetamol (Paracet®, Weifa, Oslo, Norway) as well as 10 mg oxycodone sustained release tablet (OxyContin®, Dublin, Ireland) for postoperative pain management. Propofol (Propolipid®, Fresenius Kabi, Uppsala, Sweden) and remifentanil (Ultiva®, GlaxoSmithKline, Parma, Italy) were administered by computer-controlled infusion pumps (B Braun Perfusor Space®, Melsungen, Germany) using a target-controlled infusion (TCI) program (Schnider for propofol and Minto for remifentanil) in order to achieve plasma concentrations sufficient for anesthesia and analgesia. Prior to start of anesthesia all patients received an infusion of Ringer’s-Acetate (5 ml /kg) to prevent hypotension during anesthesia induction, as well as 3 - 5 ml 1 % lidocaine intravenously to prevent pain during propofol injection. All patients were pre-oxygenated with 100 % oxygen and received the non-depolarizing muscle relaxant cisatracurium for intubation (Nimbex®, GlaxoSmithKline, Oslo, Norway). After intubation the inspiratory oxygen fraction was reduced to 40 %; nitric oxide was not used.

### Data Preprocessing

#### Study 1 - Sleep scalp EEG

Data was imported to EEGLAB(56) and epoched into 5 seconds bins. Epochs that contained artifacts (e.g. eye blinks or movement) were manually inspected and rejected by a trained scorer (B.A.M.). None of the channels were discarded or interpolated. On average, the participants had 5748.9 ± 10.01 of these five second epochs and 946.95 ± 542.68 of them were rejected (16.44 ± 2.98 %). The data from the healthy sleep participants has been published before and was cleaned in a comparable approach(26, 27, 52, 53). For further analysis in MATLAB (MATLAB Release R2017b, The MathWorks, Inc., Natick, Massachusetts, United States) the data was then imported into FieldTrip(57).

#### Study 2 – Sleep intracranial EEG

Data was imported to FieldTrip(57), downsampled to 500 Hz and segmented into 30 seconds segments for subsequent data analysis. Anatomical localization was carried out by fusing pre-implantation T1-weighted Magnetic Resonance Imaging (MRI) scans with post-implantation MRI and both automatic and manual labelling of the electrode position (see above). Epileptic, white matter and channels with other artifacts were discarded. The data was bipolar referenced, demeaned and detrended.

#### Study 3 - Anesthesia scalp EEG

Data was imported into FieldTrip(57) and epoched in 10 second bins. An Independent Component Analysis (fastica(58)) was used to clean the data from systematic artifacts such as the ECG. Further data cleaning was done manually after inspection by a neurologist (R.T.K.) and an anesthesiologist (J.D.L). On average, the patients had 1183 ± 81.42 ten second epochs of which 196 ± 103.19 were marked as noisy (15.81 ± 3.15 %); comparable to the sleep EEG study (Study 1). No channels were excluded or interpolated. Data was referenced using the common average, demeaned and detrended. Wake periods were defined as time before induction and after anesthesia when the patients responded reliably to verbal commands of the study personnel. Anesthesia periods were defined as time after induction until the termination of propofol application.

#### Study 4 – Anesthesia intracranial EEG

Data was recorded with a 512 Hz digitization rate in eight patients. Four additional patients were recorded with a digitization rate of 1024 Hz and these datasets were down-sampled to 512 Hz. Data was then imported to FieldTrip(57), epoched into 10-second segments and inspected by a neurologist (R.T.K.) for epileptic activity and manually cleaned of epileptic and other non-neural artifacts. The awake state was defined as time before start of propofol, anesthesia was defined as time after loss of consciousness (unresponsiveness to verbal commands assessed by study personnel and attending anesthetist). After fusing the pre-implantation T1-weighted MRI and the post-implantation Computer Tomography (CT) scans, electrodes were automatically localized by an openly available brain atlas (Freesurfer(59)) in parallel with manual positioning by experienced neurologists for cross validation. Contacts in white matter or lesions were discarded. The remaining signals were then bipolar referenced to their lateral neighbor, demeaned and detrended.

### Spectral Analysis

(1) To obtain average power spectra, after artifact removal the data was epoched into 10 second segments for anesthesia and 30 second segments for sleep. (2) Time-frequency decomposition was accomplished by using the Fast Fourier Transformation (*mtmfft*, FieldTrip(57)) from 0.5 Hz to 45 Hz in 0.5 Hz steps. The analysis was limited to 45 Hz due to line noise at 50 Hz in the Oslo recordings and then adopted to all consecutive studies for consistency. To obtain reliable spectral estimates we utilized a multi-taper approach based on discrete prolate slepian sequences (dpss; anesthesia: 9 tapers for 10 second segments, no overlap, frequency smoothing of ± 0.5 Hz; sleep: 29 tapers for 30 second segments, no overlap, frequency smoothing of ± 0.5 Hz). The power spectrum of each state was averaged over all samples of the state (rest or wake, non-rapid eye movement sleep stage 3 (N3) and rapid eye movement sleep (REM) or wake and anesthesia), channels and subjects (Fig. 1b, c and Fig. 2b, c). For better comparison, we visualized the effect on the scalp level. For study 4 no simultaneous EEG recordings were available. (2) To elucidate the time frequency relationship over time as depicted in Figure 1a and 2a, we again employed a multi-taper spectral analysis of frequencies between 0.5 and 45 Hz in 0.5 Hz steps this time using a sliding, overlapping time window (anesthesia: 10 seconds, 96% overlap, frequency smoothing of ± 0.5 Hz and 9 dpss tapers; sleep: 30 seconds, 85% overlap, frequency smoothing of ± 0.5 Hz and 29 dpss tapers).

### Spectral slope estimation

We calculated the spectral slope by fitting a linear regression line to the higher frequency 1/f slope of the power spectrum in the range from 30 - 45 Hz, because it had been shown that fitting in this range best correlates with the E/I balance(22). In line with previous reports, we excluded the low frequencies that contain strong oscillatory responses, which distort the linear fit as well as the range over 50 Hz, which is confounded by both line noise (50 Hz in Europe, 60 Hz in the US) as well as broad-band muscle artifacts.

We then adapted this range to the calculation of the slope in the other studies for consistency reasons. To compute a time resolved estimate of the spectral slope, we calculated the best line fit to the 10 (anesthesia) or 30 (sleep) second segments of the multi-tapered power spectra (see above) in log-log space using polynomial curve fitting (*polyfit.m*, MATLAB and Curve Fitting Toolbox Release R2015a, The MathWorks, Inc., Natick, Massachusetts, United States). One subject in Study 1 (sleep EEG) had only noisy wake trials; therefore, his data had to be excluded from all slope comparisons to wakefulness.

### Mutual Information

Mutual Information (MI) is a metric of information theory to assess the mutual dependence of the two signals, specifically the amount of information gained about one variable when observing the other(60). This is particularly useful for non-linear, binned signals that need to be analyzed independent of rank. Mutual information between the two signals X and Y is defined as

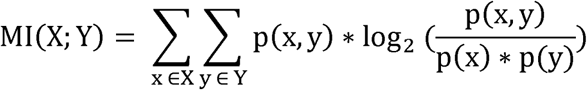

where p(x,y) depicts the joint probability function and p(x) and p(y) indicate the class probabilities. Probabilities were normalized by their sum. For MI analysis (Fig. 1b, Fig. S4, S7, S10), we epoched the time-resolved slope into 30 second segments (the hypnogram was staged in 30 second epochs) and discretized it into five bins from minimum to maximum (Wake, REM, N1, N2, N3) using the discretize.m function of MATLAB Signal Processing Toolbox Release R2015a (MathWorks Inc., USA).

### Spectral slope estimation during a slow wave

Slow wave events (Fig. S8, S9) were detected for each channel based on established algorithms(61): The raw signal was bandpass-filtered between 0.16 and 1.25 Hz and zero crossings were detected. Events were then selected using a time (0.8 to 2 s duration) and an amplitude criterion (75 % percentile). The raw data was then epoched relative to the trough of the slow wave (± 2.5 s). Time-frequency decomposition was computed in 500 ms time windows with a 250 ms overlap using FieldTrip(57) (mtmfft, frequency smoothing of ± 2 Hz and 1 dpss taper). The spectral slope was calculated by the best line fit in these time windows in log-log space between 30 - 45 Hz using polynomial curve fitting (*polyfit.m*, MATLAB and Curve Fitting Toolbox Release R2015a, MathWorks, Inc., USA).

### Classification analysis

We employed a linear discriminant analysis (LDA) to assess if slow wave power or the spectral slope were a better predictor of wakefulness or sleep. We utilized a leave-one-exemplar-out cross-validation approach that was repeated 50 times after randomly sampling an equal number of sleep and REM trials to equate the number of samples. Then every sample of the subsampled distribution was held out of the training dataset once. The LDA classifier was trained on the remaining samples and tested on the held-out test sample. The classifier performance was then assessed as percent correct. Two of the 20 sleep EEG participants had to be excluded due to insufficient number of wake trials.

### Statistical testing

To compare three states (awake, NREM and REM), we utilized Greenhouse-Geisser corrected 1-way repeated measures analysis of variance (Fig. 1b, 1c; RM-ANOVA). Effect size was calculated using Cohen’s d. The spectral slope of the awake and anesthetized state was compared using Student’s t-test for paired samples (Fig. 2b, c).

To assess the spatial extent of the observed effects in EEG, we calculated cluster-based permutation tests to correct for multiple comparisons as implemented in FieldTrip(57) (Monte-Carlo method; maxsize criterion; 1000 iterations). A permutation distribution was obtained by randomly shuffling condition labels and then compared to the actual distribution to obtain an estimate of significance. Spatial clusters are formed by thresholding independent t-tests of slope differences between wake and sleep (Fig. 1b) or wake and anesthesia (Fig. 2b) at a p value < 0.05. All results were Bonferroni-corrected for multiple comparisons. In order to control for EMG as a potential confound in the sleep EEG (Study 1), we utilized a partial correlation (Spearman) that partialled the slope of the EMG out of the correlation before computing the cluster-based permutation test (Fig. S4a). Correlation coefficients (r-values) were transformed into t-values using the following formula (N = number of subjects):

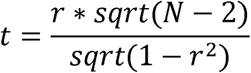

For statistical assessment of the Mutual Information, we employed surrogate testing (Fig. 1b, Fig. S4a). To obtain a surrogate distribution from the observed data, we utilized a random block swapping procedure(62, 63). The number of repetitions was equal to the number of available sleep stages. On every iteration, we re-calculated the MI of these block swapped hypnograms with the discretized time-resolved slope to create a surrogate distribution against which we could compare our original observation. To compare the results across subjects, we z-scored the values by subtracting the mean of the surrogate distribution from the observed MI and dividing by the standard deviation of the surrogate distribution. Note that a z = 1.96 reflects an uncorrected two-tailed p-value of 0.05, while a z-score of >2.8 indicates a Bonferroni-corrected significant p-value (p < 0.05 / 19 channels = 0.0026). The z-values were transformed into p-values for topographic display (Fig.1b; Fig. S4a) based on a normal cumulative distribution function (two-tailed).

### Connectivity

For the analysis of fronto-parietal connectivity (Fig. S10), we choose electrode Fz and Pz in our sleep EEG recordings (Study 1; n = 20) to calculate the magnitude squared coherence from frequencies of 0.1 to 64 Hz in 0.1 Hz steps using the mscohere.m function from the MATLAB Signal Processing Toolbox and described previously(39, 40). Note that coherence estimates reflect both power changes as well as changes in phase synchrony. Therefore, we also calculated the Phase-Locking Value (PLV) and amplitude correlations (rho) to disentangle the effects of phase and power, respectively. To discount the effects of volume spread, we calculated the imaginary PLV(64) (iPLV) and orthogonalized power correlations(65) (rhoortho).

We then quantified the Mutual Information(60) (MI; see above) to compare how well the results capture the changes between different sleep stages across the night. For this analysis we only utilized the slope values of electrode Fz (as we were calculating the other measures in Fz-Pz) and defined theta from 4-10 Hz analog to Pal et al.(39, 40).

## Supplemental Figures

**Fig. S1:**
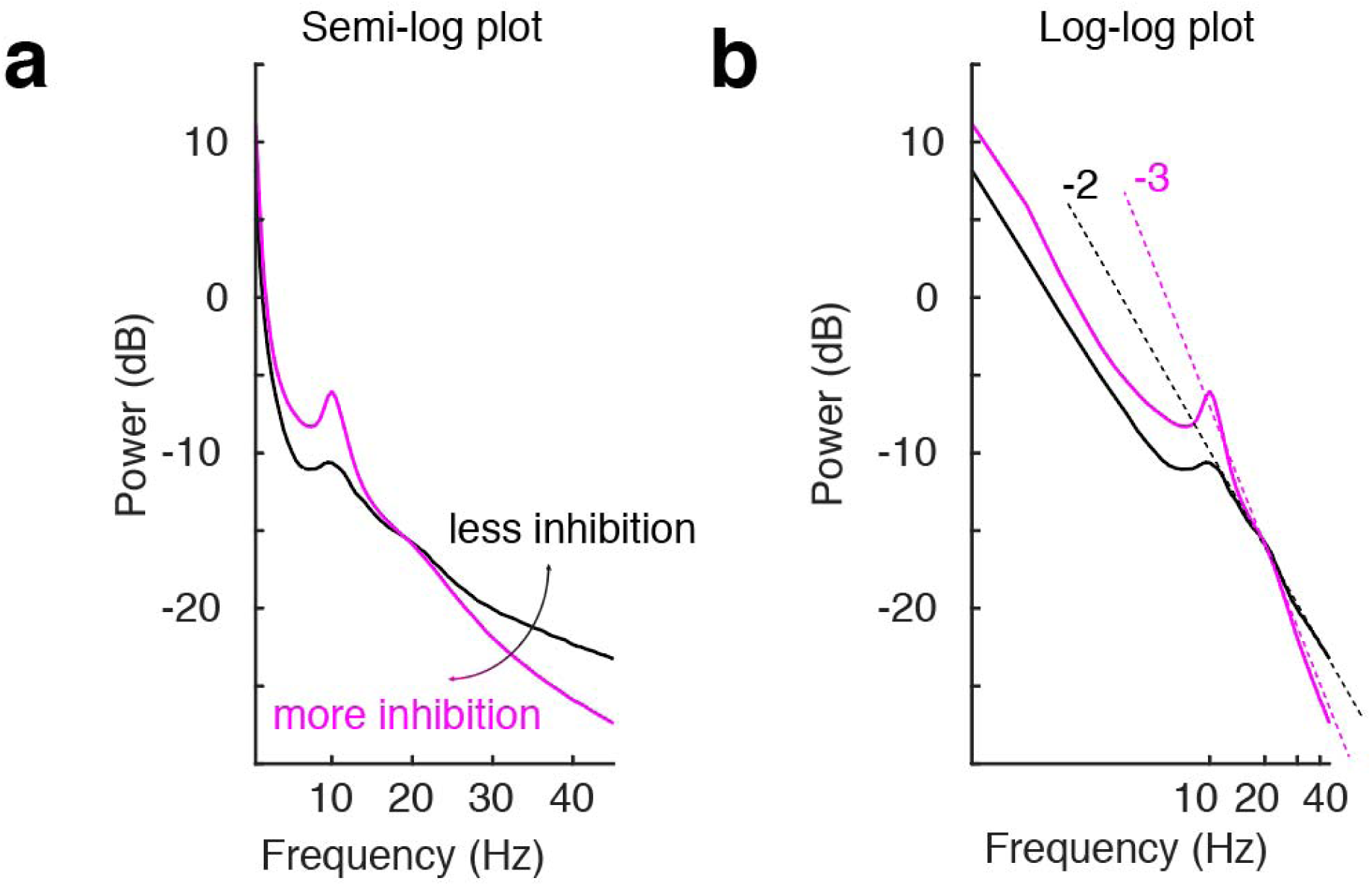
The spectral slope - a surrogate marker for excitation / inhibition balance. **a**, Power spectral density (PSD) in semi-log plot. Example wake (black) and anesthesia PSD (magenta). More inhibition results in a steeper decrease of the PSD in frequencies above 30 Hz. **b**, PSD in a log-log plot. Example wake (black) and anesthesia PSD (magenta) with linear fits to 30 – 50 Hz for both states (dotted lines). The linear fit reveals a more negative spectral slope for states with higher inhibition.

**Fig. S2:**
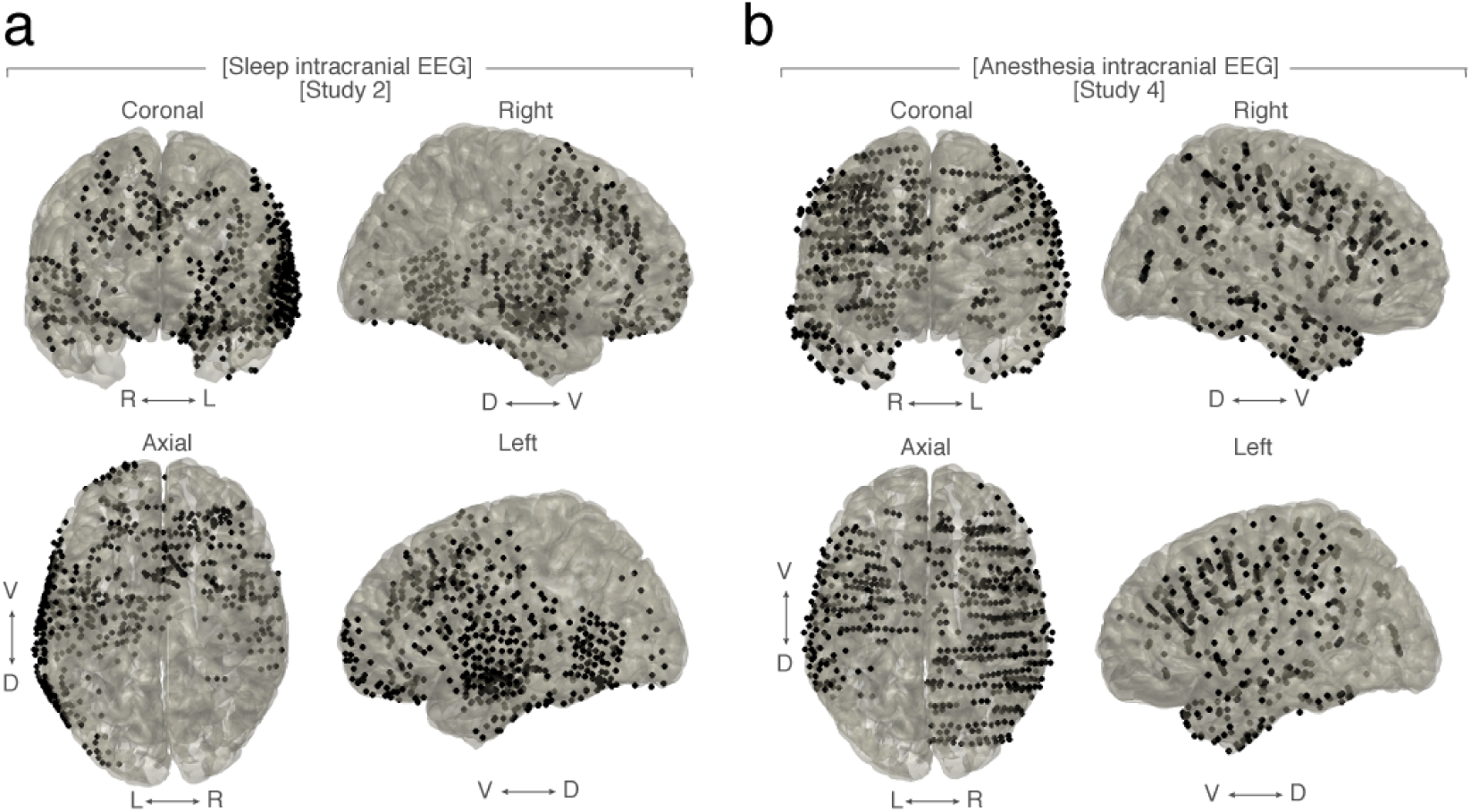
Coverage in intracranial subjects. **a,** Sleep intracranial EEG – Grid and SEEG contacts of all subjects (n = 10) plotted on MNI brain. Right (R), left (L), ventral (V), dorsal (D). **b**, Anesthesia intracranial EEG – Grid, Strip and SEEG contacts of all subjects (n = 12) plotted on a Montreal Neurological Institute (MNI) brain. Right (R), left (L), ventral (V), dorsal (D).

**Fig. S3:**
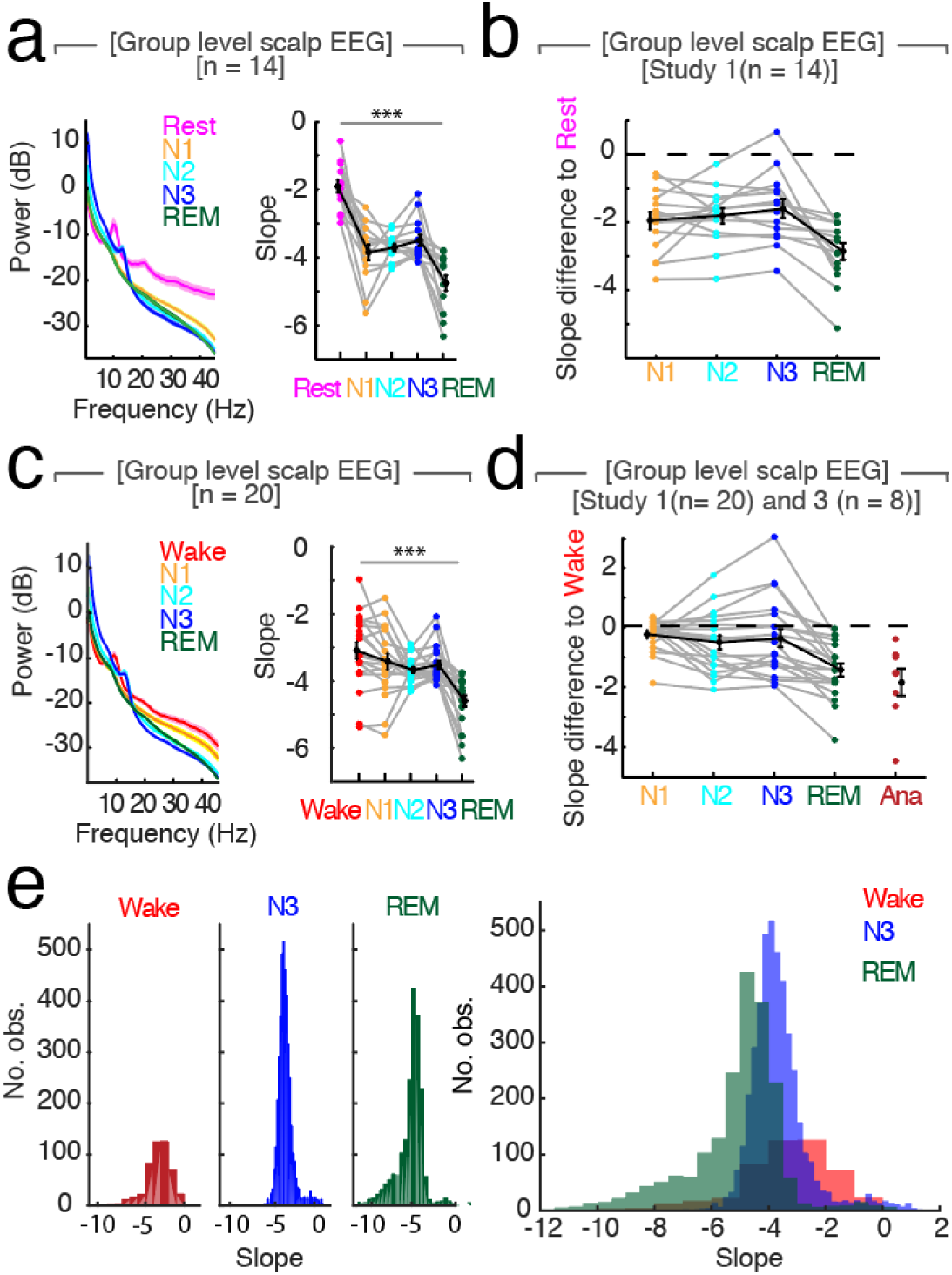
Relative changes of spectral slope reliably differentiate between wakefulness, sleep and general anesthesia. **a**, Left – Mean power spectra (± SEM) averaged across all channels and subjects (n = 14) during rest recording of 5 min eyes closed recorded before sleep compared to all sleep stages. Right – Slope values. Mean ± SEM in black. Repeated measures ANOVA: *** p < 0.001, F_2.54, 33.02_ = 38.02, d_Rest-Sleep_ = 3.52. Post-hoc t-tests: p_Rest-N2_ < 0.001; t_13_ = 7.97; d = 3.31; p_Rest-N3_ < 0.001; t_13_ = 5.69; d = 2.49; p_Rest-REM_ < 0.001; t_13_ = 11.67; d = 3.71; p_N3-REM_ < 0.0001; t_13_ = 4.44; d = 1.70. **b**, Slope differences of all sleep stages to rest (n = 14). Mean ± SEM in black. **c**, Left - Mean power spectra (± SEM) averaged across all channel and subjects (n = 20) during wakefulness and all sleep stages. Right - Slope values. Mean ± SEM in black. Repeated measures ANOVA: *** p < 0.001, F_1.86, 33.49_ = 13.39, d_Wake-Sleep_ = 0.79. Post-hoc t-tests: p_Wake-N2_ = 0.029, t_18_ = 2.36, d = 0.69; p_Wake-N3_ = 0.19; t_18_ = 1.34; d = 0.48; p_Wake-REM_ < 0.0001; t_18_ = 6.83; d = 1.58; p_N3-REM_ < 0.0001; t_19_ = 5.12; d = 1.66. **d**, Slope difference of all sleep stages to all wake trials (n = 20) and anesthesia to wake trials before anesthesia (n = 8). Mean ± SEM in black. **e**, Histogram of slope values pooled across all participants (n= 20). Wakefulness (red), N3 (blue), REM (green). Left: Separated values of each sleep stage. Right: All three sleep stages within one plot.

**Fig. S4:**
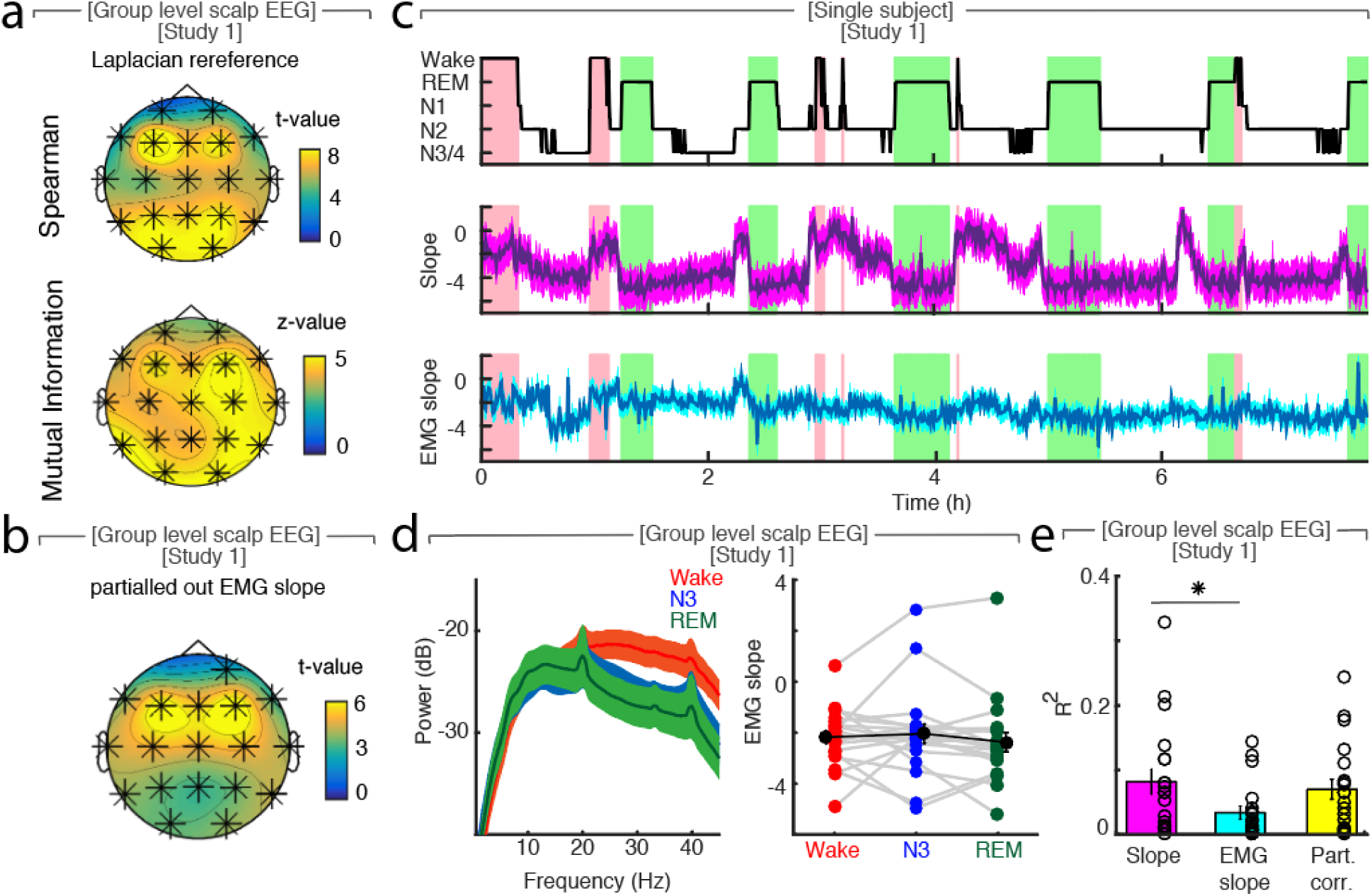
The spectral slope is not confounded by muscle activity. **a**, Laplacian re-reference (n = 20). Upper topoplot: Cluster permutation test of Spearman rank correlation between hypnogram and time-resolved slope. * p< 0.05. Lower topoplot: Mututal Information between slope and hypnogram. Statistic with random block swapping. * p < 0.05. **b**, Cluster permutation test of Spearman rank correlation between hypnogram and time-resolved slope with electromyography (EMG) slope partialled out (n = 20). * p < 0.05. **c**, Hypnogram of a single subject (upper panel), time-resolved slope averaged over EEG channels F3, Fz and F4 (middle panel), time-resolved slope of EMG signal averaged over three EMG channels (lower panel). **d**, EMG signal on group level across a full night (n = 20). Left: Power spectra of EMG (mean ± SEM). Right: Slope of EMG in wakefulness (red), NREM stage 3 (blue), REM (green), grand average (black, mean ± SEM). **e**, R^2^ of Spearman rank correlations averaged across all channels between hypnogram and slope (magenta), EMG slope (cyan) and the slope with the EMG slope partialled out (yellow, all mean ± SEM). The correlation of hypnogram - slope and hypnogram - EMG slope is significantly different (paired t-test: p = 0.0059, t_19_= 3.10). Furthermore, we utilized the LDA classification approach to test if the spectral slope outperforms the EMG slope for state discrimination. We found that the spectral slope performed significantly better at distinguishing all three states (t = 4.19, p < 0.001, d = 1.24; slope: 58.09 ± 2.35%, EMG slope: 46.03 ± 2.12%; chance 33%). Likewise, the slope was better at discriminating only WAKE and REM (t = 3.03, p = 0.008, d = 0.89; slope: 76.32 ± 3.61%, EMG slope: 64.89 ± 2.24%; chance 50%).

**Fig. S5:**
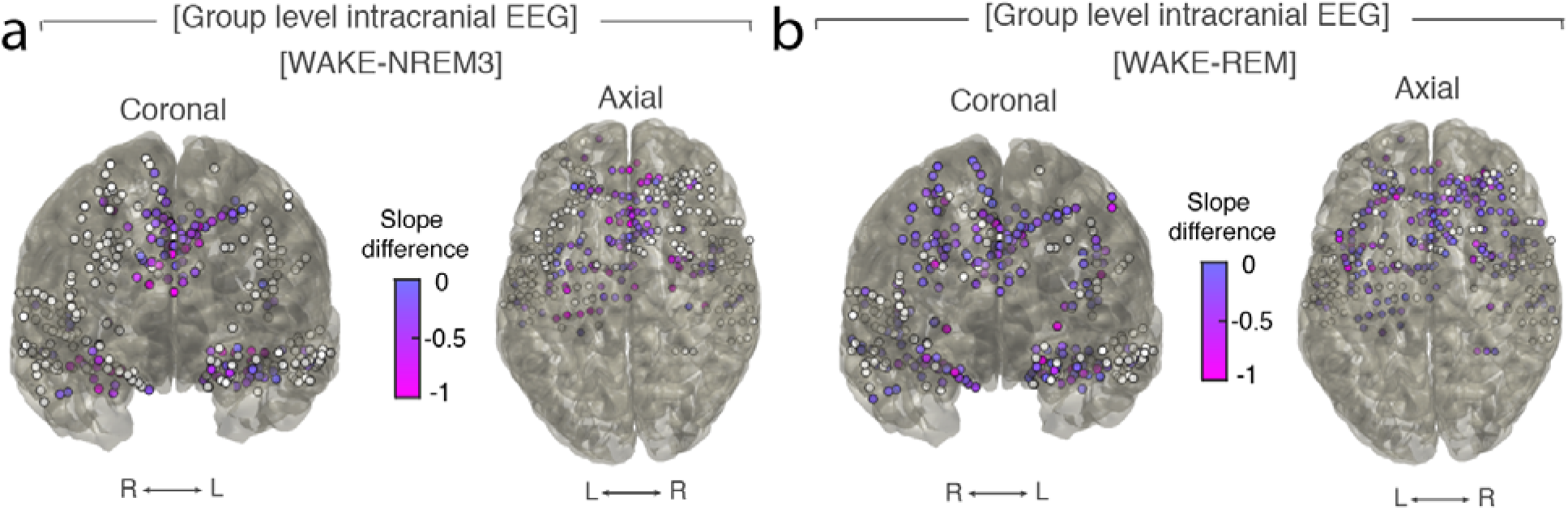
Differences of spectral slope in intracranial electrodes between waking and NREM 3 or REM sleep (n = 10). **a**, Left – coronal, right – axial view of electrodes that followed observed EEG pattern with a more negative slope for NREM 3 sleep than for waking (magenta). Electrodes that did not show the pattern are depicted in white. Right (R), left (L). **b**, Left – coronal, right – axial view of electrodes that followed observed EEG pattern with a more negative slope for REM sleep than for waking (magenta). Electrodes that did not show the pattern are depicted in white. Right (R), left (L).

**Fig. S6:**
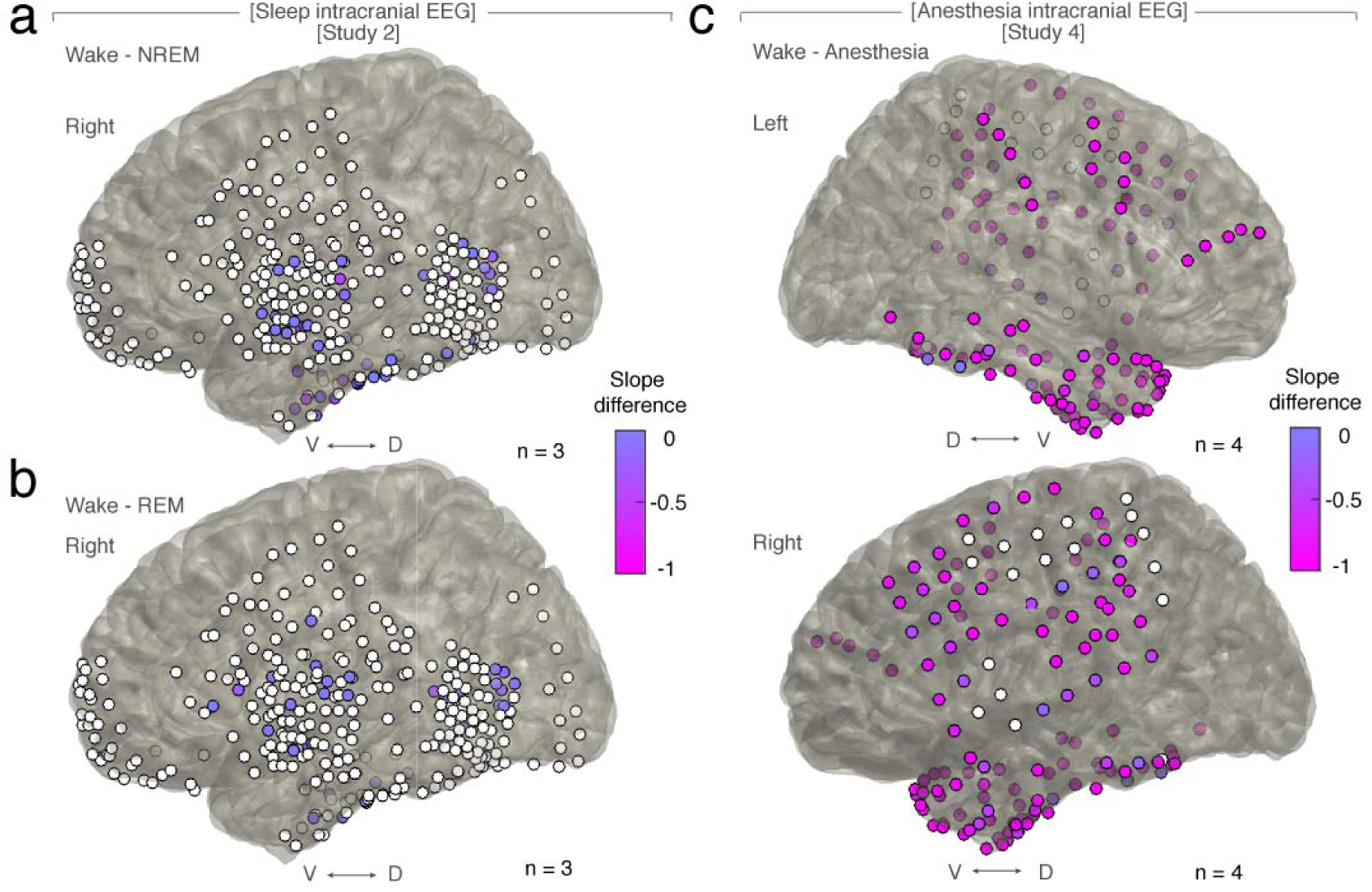
Differences in spectral slope in sleep and under general anesthesia in cortical electrodes. **a**, All grid and strip contacts of 3 patients plotted on MNI brain. Electrodes that followed the pattern of more negative slope in NREM stage 3 sleep than in waking are colored purple to magenta. Electrodes that did not show the pattern are depicted in white. Ventral (V), dorsal (D). **b**, All grid and strip contacts of 3 patients plotted on MNI brain. Electrodes that followed the pattern of more negative slope in REM sleep than in waking are colored in purple to magenta. Electrodes that did not show the pattern are depicted in white. Ventral (V), dorsal (D). **c**, All grid and strip contacts of 4 patients plotted on MNI brain. Electrodes that followed the pattern of more negative slope under anesthesia than in waking are colored in purple to magenta. Electrodes that did not show the pattern are depicted in white. Right (R), left (L), ventral (V), dorsal (D).

**Fig. S7:**
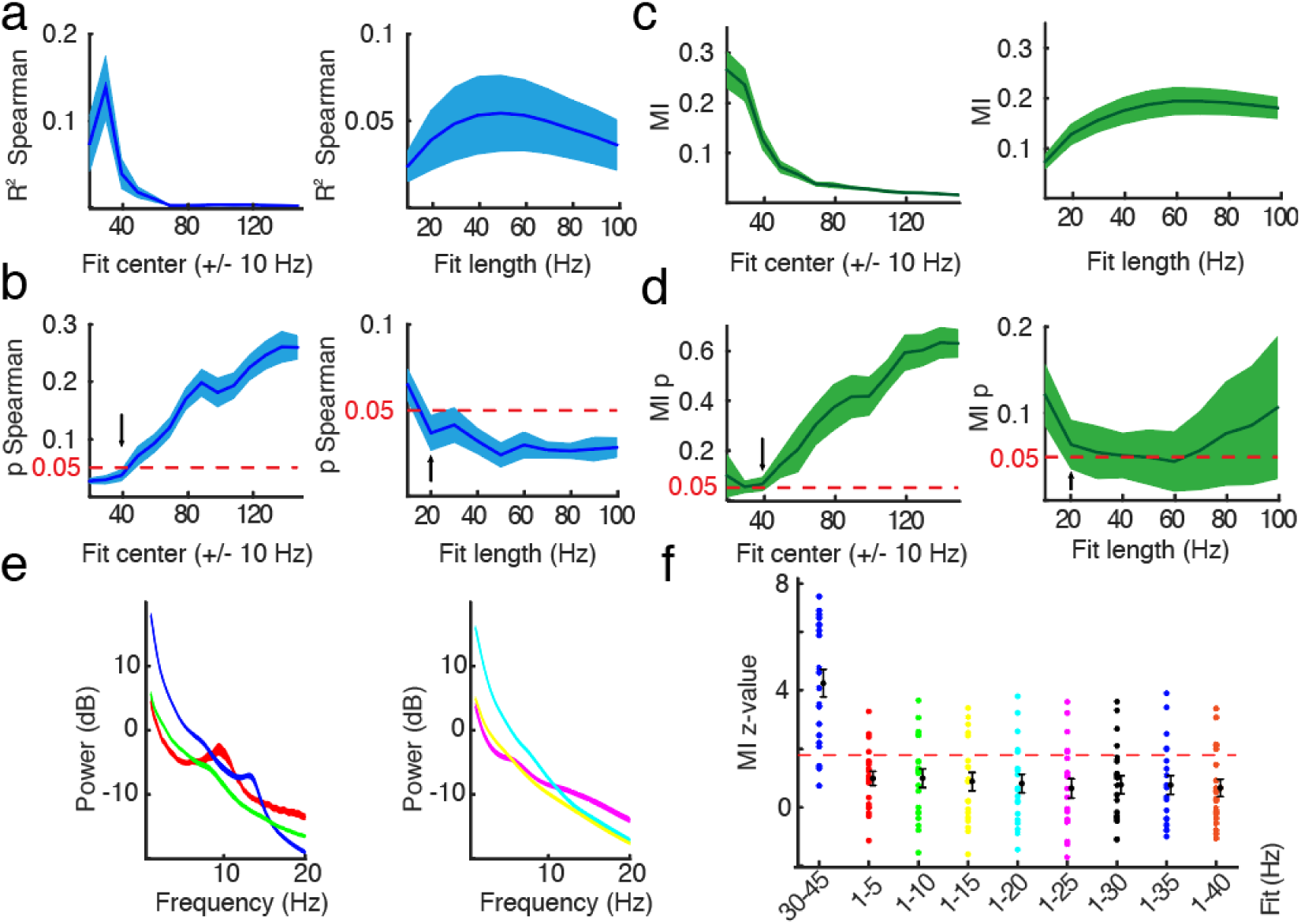
Evaluation of different slope fit settings in intracranial sleep. **a**, Spearman rank correlation (R^2^) between slope and hypnogram with different slope fits with center frequencies from 20 to 150 ± 10 Hz with SEM in intracranial data during sleep (blue; n = 10). Red dotted line for p value of 0.05. Black arrow indicates used center frequency of 40 Hz (30 – 50 Hz) for this study. **b**, Spearman rank correlation (R^2^) with different slope fit length from 30 to 40 Hz up to 30 to 130 Hz (10 – 100 Hz fit length) with SEM (blue). Red dotted line for p value of 0.05. Black arrow indicates used fit length for this study (20 Hz; 30 – 50 Hz). To control for the shared variance between SO power and the spectral slope, we repeated the correlations and partialled out the corresponding SO power, which left the results unchanged (t_19_ = 1.37, p = 0.188, d = 0.21; before: R^2^ = 0.13 ± 0.03; after: R^2^ = 0.09 ± 0.03). **c**, Mutual Information (MI) between slope and hypnogram with different slope fits with center frequencies from 20 to 150 ± 10 Hz with SEM in intracranial data during sleep (green; n = 10). Red dotted line for p value of 0.05. Black arrow indicates used center frequency of 40 Hz (30 – 50 Hz) for this study. **b**, Mutual Information (MI) with different slope fit length from 30 to 40 Hz up to 30 to 130 Hz (10 – 100 Hz fit length) with SEM (green). Red dotted line for p value of 0.05. Black arrow indicates used fit length for this study (20 Hz; 30 – 50 Hz). **e**, Mixed (left) and fractal component (right) of power spectra in scalp EEG (n = 20) after IRASA. **f**, Z-value of surrogate distribution (random block swapping) of Mutual Information (MI) between slope and hypnogram using the original (blue, 30-45 Hz) and different slope fits to fractal component (obtained by IRASA) in lower frequencies. Note that a z = 1.96 reflects an uncorrected two-tailed p-value of 0.05, while a z-score of >2.8 indicates a Bonferroni-corrected significant p-value (p < 0.05 / 19 channels = 0.0026).

**Fig. S8:**
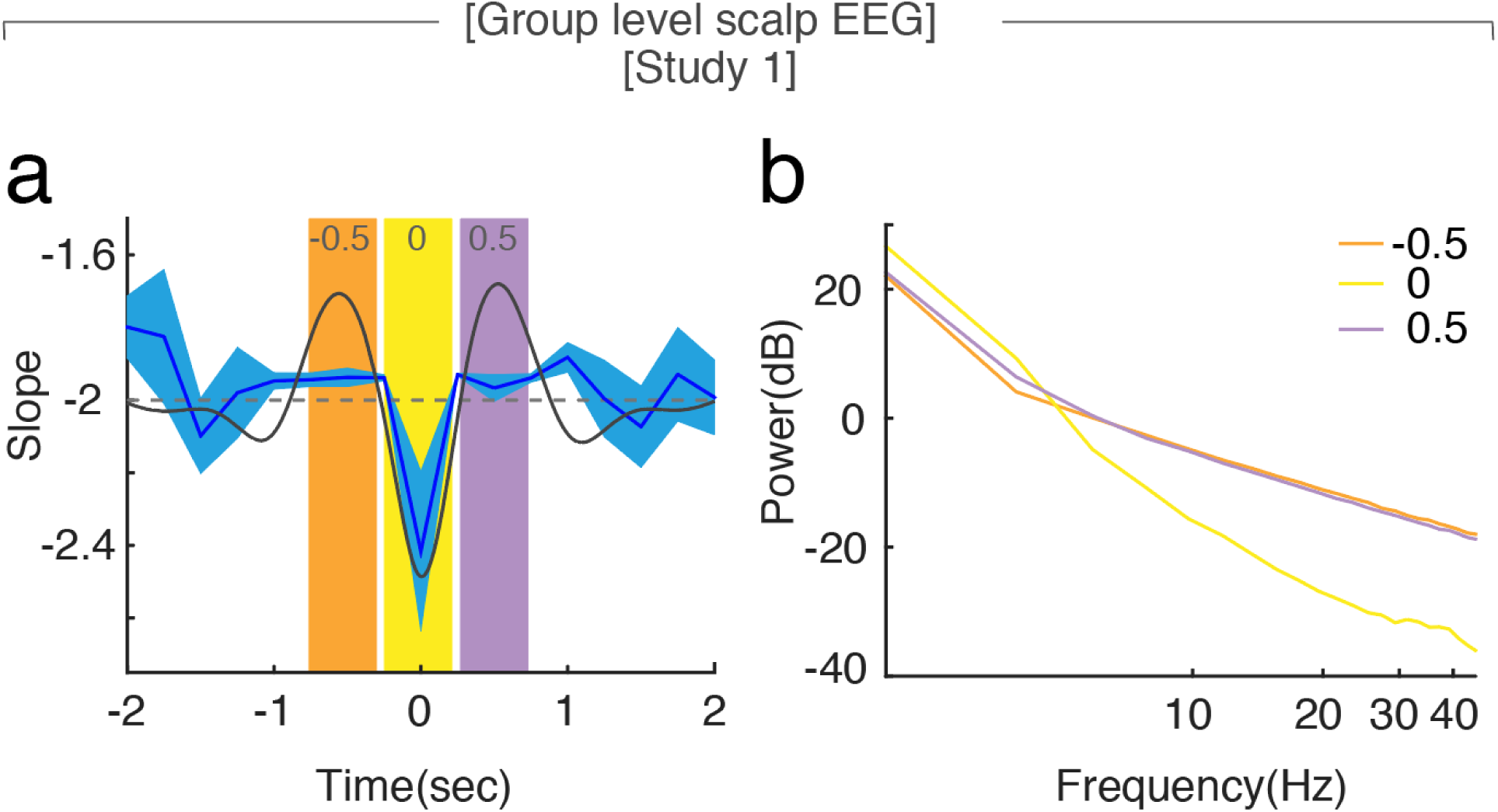
The spectral slope tracks changes in inhibition during slow waves. **a**, Average spectral slope changes over the time course of all slow waves in scalp EEG (n = 20) during sleep (blue; mean ± SEM), Superimposed in gray is the average slow wave of all subjects. Highlighted are the following time windows: −750 to −250 (orange), −250 to 250 (yellow) and 250 to 500 ms (purple). **b**, Power spectra in log-log space within specified time windows during the slow wave: −750 to −250 (center: −0.5 s; orange), −250 to 250 (center: 0 s; yellow) and 250 to 500 ms (center: 0.5 s; purple). Note the steep power decrease during the through of the slow wave (yellow).

**Fig. S9:**
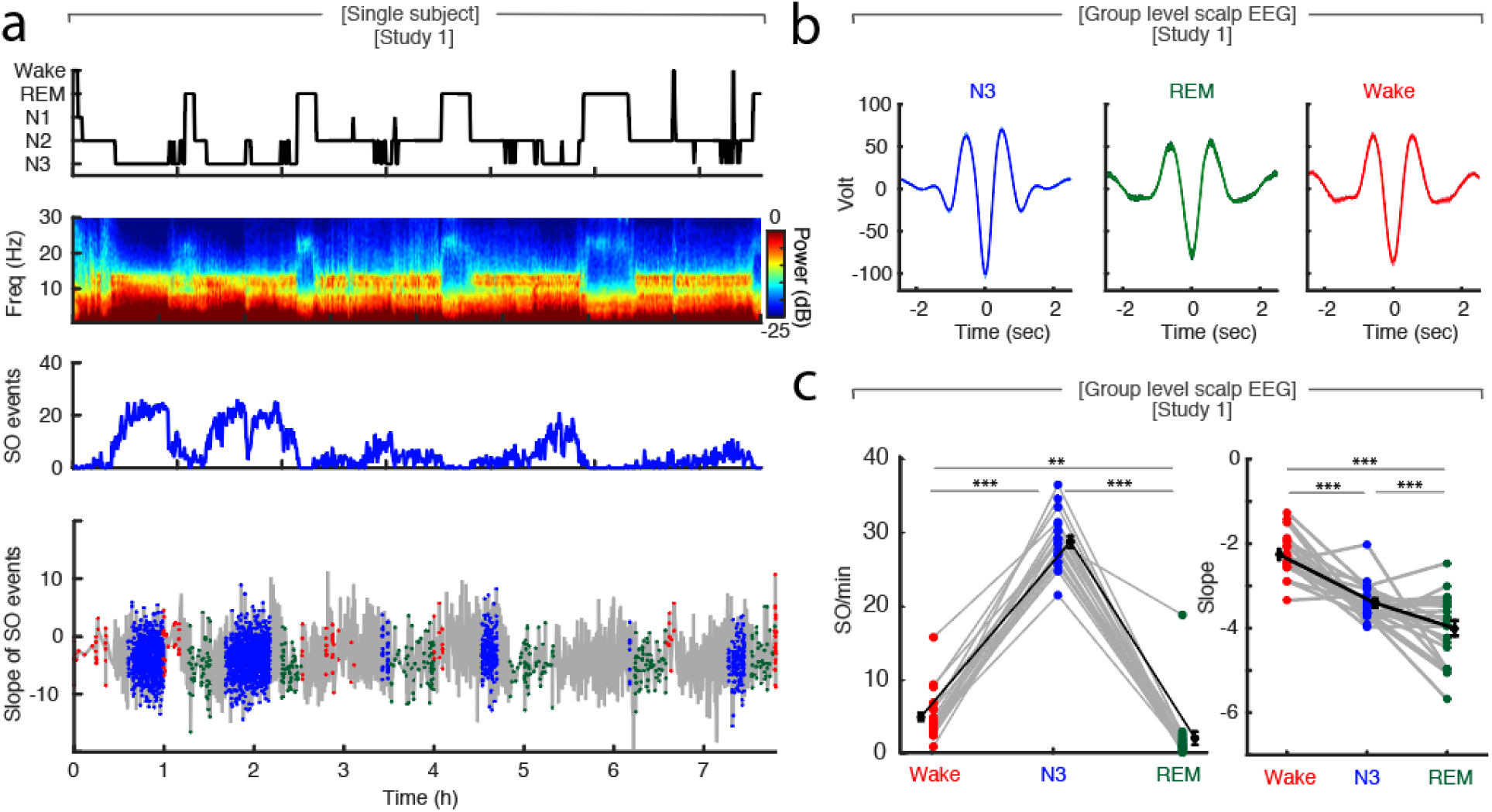
Slow waves during wakefulness, N3 and REM sleep in scalp EEG. **a**, Single subject example: Upper panel: Hypnogram. Upper middle panel: Multitapered spectrogram of electrode Fz. Lower middle panel: Number of slow wave (SO) events during 30 second segments of sleep in electrode Fz. Note the decreasing number of SO events during the course of the night. Lower panel: Spectral slope of SO events occurring in N3 (blue), wakefulness (red) and REM sleep (green) in electrode Fz. Background: Time-resolved slope of electrode Fz in light grey. **b**, Group level (n = 20) average waveforms in electrode Fz during N3 (blue), REM sleep (green) and wakefulness (red; mean ± SEM). **c**, Left: Slow wave events per minute in wakefulness (red), N3 (blue) and REM (green) in scalp EEG channel FZ (n = 20). In black mean ± SEM. Paired t-test: ** p < 0.01, *** p< 0.001. Right: Slope of slow wave events on the group level (n = 20; averaged across all 19 EEG electrodes) in wakefulness (red), N3 (blue) and REM sleep (green). Mean ± SEM in black. Paired t-test: *** p< 0.001.

**Fig. S10:**
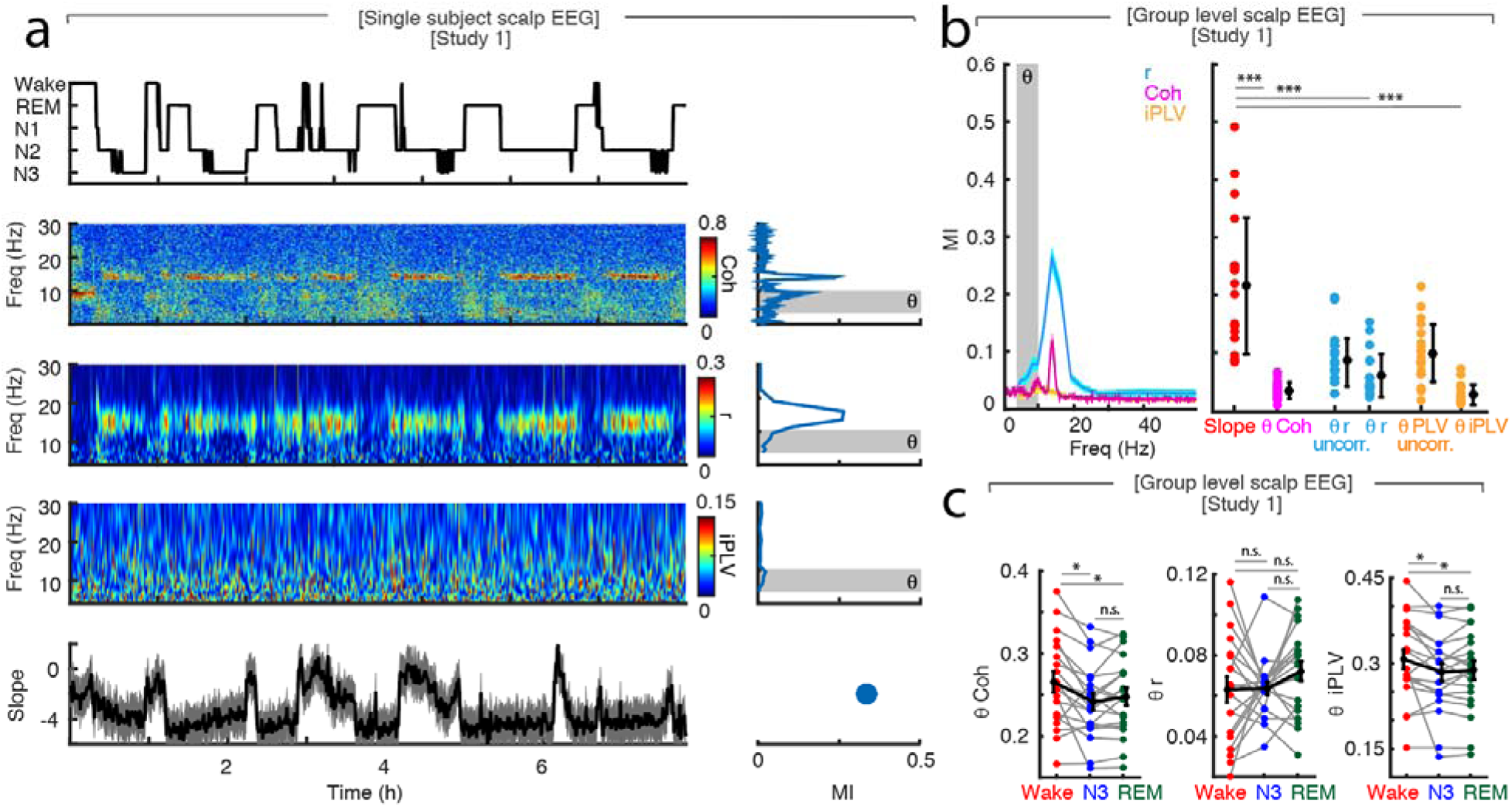
Comparison of Mutual Information captured by fronto-parietal connectivity and spectral slope. **a,** Single subject example: Upper panel – hypnogram. Middle panels – Fz-Pz coherence (Coh), Fz-Pz connectivity measured by orthogonalized power correlation (r) and imaginary phase-locking value (iPLV) between 0.1 and 30 Hz. Right subpanels - Accompanying mutual information between the hypnogram and all frequencies, theta (θ; 4-10 Hz) highlighted in grey. Lower panel – spectral slope (30 - 45 Hz) of Fz. Right subpanel – Mutual information of the Fz slope. **b,** Group level (n = 20) analysis of mutual information for Fz-Pz coherence and connectivity measured by orthogonalized power correlation (r) or imaginary phase-locking value (iPLV). Left panel – Across all frequencies. Right panel – Comparison of mutual information between Fz slope, theta (θ) coherence and connectivity measured by power correlation (uncorrected and orthogonalized) as well as phase-locking value (uncorrected and imaginary). Paired t-test: *** < 0.001. **c**, Group level (n = 20) comparison of Fz-Pz theta (θ) coherence, orthogonalized power correlation (r) and weighted phase-locking value (iPLV) between wakefulness, N3 and REM sleep, showing that these metrics do not reliably distinguish between N3/SWS and REM sleep. Paired t-test: n.s. – not significant, * p<0.05.

**Fig. S11:**
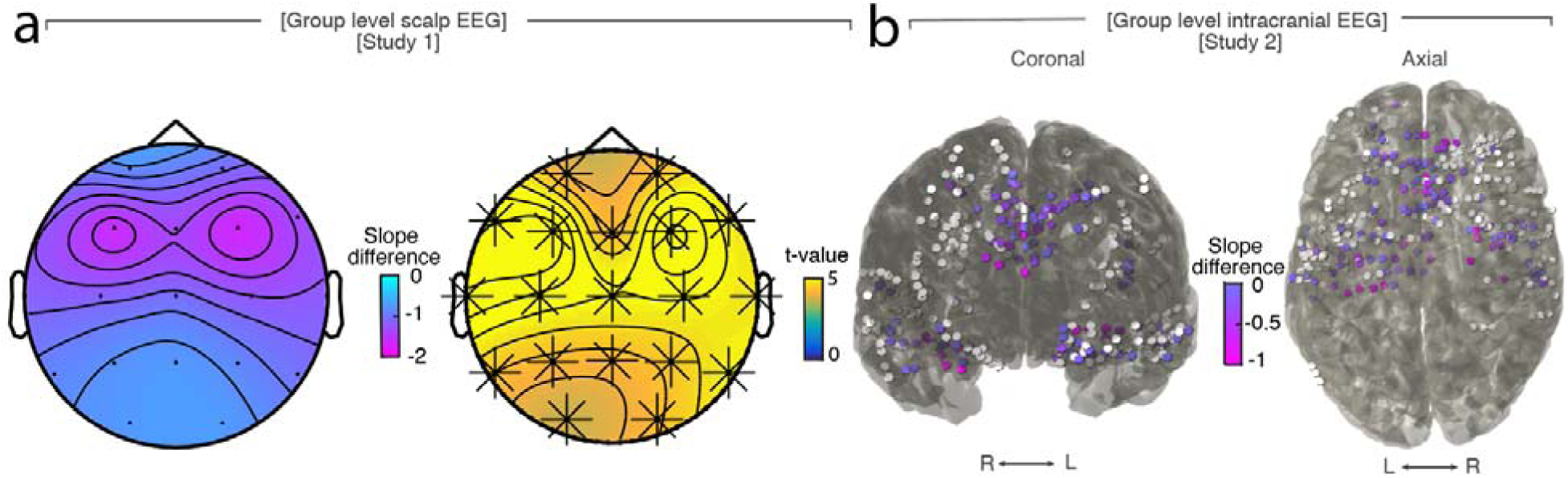
Slope difference between N3 and REM sleep. **a**, Scalp EEG (n = 20). Left: Topography of slope difference, Right: Cluster permutation test between slope of N3 and REM. * p < 0.05. **b**, Depth electrodes (n = 10). Left – Coronal view. Right – Axial view on an MNI brain contain all intracranial electrodes of all patients. Colored – contacts that showed a more negative slope in REM compared to N3 slope. White – contacts that did not show the pattern.

